# Using Polysome Isolation with Mechanism Alteration to Uncover Transcriptional and Translational Dynamics in Key Genes

**DOI:** 10.1101/006213

**Authors:** Bradly Alicea, Steven T. Suhr, Jose B. Cibelli

## Abstract

What does it mean when we say a cell’s biochemistry is regulated during changes to the phenotype? While there are a plethora of potential mechanisms and contributions to the final outcome, a more tractable approach is to examine the dynamics of mRNA. This way, we can assess the contributions of both known and unknown decay and aggregation processes for maintaining levels of gene product on a gene-by-gene basis. In this extended protocol, drug treatments that target specific cellular functions (termed mechanism disruption) can be used in tandem with mRNA extraction from the polysome to look at the dynamics of mRNA levels associated with transcription and translation at multiple stages during a physiological perturbation. This is accomplished through validating the polysome isolation method in human cells and comparing fractions of mRNA for each experimental treatment at multiple points in time. First, three different drug treatments corresponding to the arrest of various cellular processes are administered to populations of human cells. For each treatment, the transcriptome and translatome are compared directly at different time points by assaying both cell-type specific and non-specific genes. There are two findings of note. First, extraction of mRNA from the polysome and comparison with the transcriptome can yield interesting information about the regulation of cellular mRNA during a functional challenge to the cell. In addition, the conventional application of such drugs to assess mRNA decay is an incomplete picture of how severely challenged or senescent cells regulate mRNA in response. This extended protocol demonstrates how the gene- and process-variable degradation of mRNA might ultimately require investigations into the course-grained dynamics of cellular mRNA, from transcription to ribosome.

## Introduction

The biochemical environment of a cell can reveal much information about changes in its phenotype (for example, see [1]). Due to the relative efficiency and accuracy of the translation and transcription [2], capturing the dynamics of mRNA molecules associated with these processes can provide a window into changes associated with cell morphology, protein expression, and other outcomes [3, 4, 5]. The artificial manipulation of cellular mRNA by exposing a cell population to specific compounds can yield insight into changes that represent shifts in gene regulation due to aging or environmental stress. In this paper, we will simulate conditions of mass arrest of cell cycle, protein synthesis, and polysomal degradation with a previously described method of assaying polysome-associated mRNA to build a model of mRNA decay and regulation in adult human fibroblast populations.

1. Jarosz, D.F., Taipale, M., and Lindquist, S. (2010). Protein Homeostasis and the Phenotypic Manifestation of Genetic Diversity: principles and mechanisms. Ann Rev Genetics, 44, 189-216.
2. Kirkwood, T.B.L., Rosenberger, R.F., and Galas, D.J. (1986). Accuracy in Molecular Processes: its control and relevance to living systems. Chapman and Hall, New York. Chapters 6 and 7.
3. Rabani, M. et.al (2011). Metabolic labeling of RNA uncovers principles of RNA production and degradation dynamics in mammalian cells. Nat Biotech, 29(5), 436-442.
4. Chechik, G. and Koller, D. (2009). Timing of gene expression responses to environmental changes. J Computational Biol, 16, 279–290.
5. Barenco, M. et.al (2009). Dissection of a complex transcriptional response using genome-wide transcriptional modelling. Mol Sys Biol, 5, 327.

## Recovery and Definition of Fractionated mRNA Pools

Rather than using high-throughput techniques, an alternate way to get at the biochemical diversity of cellular information processing is to compare translatome (abbreviated as TLT) and transcriptome (abbreviated as TST) for a range of candidate genes. While TLT measurements reveal mRNA that is loosely associated with the polysome, TST measurements reveal mRNA associated with transcription. The difference between these two fractions mainly involves the effects of mRNA decay and post-transcriptional changes. In terms of understanding biological complexity, the TLT is a critical link between TST and proteome [1, 2]. It is our contention that TLT is more directly comparable to TST than the proteome [3]. The translating ribosome affinity purification (TRAP) approach described in vivo by Heiman et.al [4] and Doyle et.al [5] allows for freshly transcribed gene products, contributing directly to protein production and precipitated straight from the polysome, to be recovered from murine CNS cells. Other studies have used similar methods to examine TLT in biological contexts ranging from cardiomyocytes [6] to stress response in yeast [7] and surveys of discrete cellular populations in *Arabidopsis* [8]. In Markou et.al [6], 5’ terminal oligopyrimidine tracts (TOPs) are used to indirectly measure translation levels and comparatively assess the activity of TLT and TST among cardiomyocyte populations. Halbeisen and Gerber [7] observe that TLT of yeast, harvested using polyribosome precipitation and bulk purification of the associated mRNA, acts to coordinate messages observed in TST during stress response. Most directly comparable with our method, Mustroph et.al [8] used polyribosome precipitation to isolate the L18 element from *in vivo Arabidopsis* specimens. They observed general enrichment of TLT during hypoxia and mosaic expression of TLT mRNA across tissue types.

The approach we use involves polysome-associated mRNA (see Methods for protocol), which is different from the transcriptional profiling approaches of Arava [9] and Ingolia [10, 11]. Approaches to transcriptional profiling focus on the small sequences of translation-associated mRNA bound to the ribosome at any given time. While transcriptional profiling can detect highly upregulated genes for certain processes (e.g. starvation in yeast) better than recovery of mRNA from bulk polysome [11], such approaches typically ask a different set of questions with regard to the data. Much of the literature on ribosomal profiling has focused on transcriptional elongation, initiation, and correlations with protein abundance [12, 13]. This generally yields smaller (<200 nt) informative sequences that provide insight into protein production and translational rate as they relate to stress, apoptosis, and cancer [13, 14]. By contrast, our approach [15, 16] yields larger informative sequences that can be understood in a quantification context common to transcriptional mRNA. This common analytical context may provide novel and potentially critical information about many types of biological processes.

1. Dressaire, C. et.al (2009). Translatome and proteome exploration to model translation efficiency and protein stability in *Lactococcus lactis*. PLoS Comput Biol, 5(12), e1000606.
2. Chang, W.Y. and Stanford, W.L. (2008). Translational control: a new dimension in embryonic stem cell network analysis. Cell Stem Cell, 2, 410–412.
3. Kettman, J.R., Frey, J.R., and Lefkovits, I. (2001). Proteome, TST, and genome: top down or bottom up analysis? Biomol Eng, 18(5), 207-212.
4. Heiman, M. et.al (2008). A translational profiling approach for the molecular characterization of CNS cell types. Cell, 135, 738–748.
5. Doyle, J.P. et.al (2008). Application of a translational profiling approach for the comparative analysis of CNS cell types. Cell, 135, 749-762.
6. Markou, T et.al (2010). Regulation of the cardiomyocyte TST vs. TLT by endothelin-1 and insulin: translational regulation of 5’ terminal oligopyrimidine tract (TOP) mRNAs by insulin. BMC Genomics, 11, 343.
7. Halbeisen, R.E. and Gerber, A.P. (2009). Stress-dependent coordination of TST and TLT in yeast. PLoS Biol, 7(5), e1000105.
8. Mustroph, A. et.al (2009). Profiling TLTs of discrete cell populations resolves altered cellular priorities during hypoxia in *Arabidopsis*. PNAS, 106(44), 18843–18848.
9. Arava, Y. et.al (2003). Genome-wide analysis of mRNA translation profiles in *Saccharomyces cerevisiae*. PNAS, 100(7), 3889-3894.
10. Ingolia, N.T. et.al (2009). Genome-wide analysis in vivo of translation with nucleotide resolution using ribosome profiling. Science, 324, 218-223.
11. Ingolia, N.T. et.al (2010). Genome-wide translational profiling by ribosome footprinting. Methods in Enzymology, 470, 119-140.
12. Reuveni, S. (2011). Genome-scale analysis of translation elongation with a ribosome flow model. PLoS Comput Biol, 7(9), e1002127.
13. Holcik, M. and Sonenberg, N. (2005). Translational control in stress and apoptosis. Nature Reviews Mol Cell Biol, 6(4), 318-327.
14. Smirnova, J.B. et.al (2005). Global gene expression profiling reveals widespread yet distinctive translational repsonses to different Eukaryotic translation initiation factors 2B-targeting stress pathways. Mol Cell Biol, 25(21), 9340-9349.
15. Angenstein, F. et.al (2005). Proteomic Characterization of Messenger Ribonucleoprotein Complexes Bound to Nontranslated or Translated Poly(A) mRNAs in the Rat Cerebral Cortex. J Biol Chem, 280(8), 6496-6503.
16. Angenstein, F. et.al (2009). A Receptor for Activated C Kinase Is Part of Messenger Ribonucleoprotein Complexes Associated with PolyA-mRNAs in Neurons. J Neurosci, 22(20), 8827-8837.

## Induction of mechanism disruption

Mechanism alteration can be defined as disruption of a key cellular process that does not result in immediate cell death. Using several forms of mechanism alteration (cell cycle, protein synthesis, and ribosomal degradation), we will identify ways to better understand the dynamics of cellular information processing during an induced biological process. To do this, we will use TLT and TST in tandem as heuristic indicators of regulatory events. Our working hypothesis is that mechanism alteration initiated by treatment with drug compounds will reveal adaptive responses in both TLT and TST. These data can likewise be modeled to demonstrate regulatory features that lie between transcriptional and translation-associated mRNA such as feedback and delays. The resulting data can also provide insight into how genes behave during transformation, and may lead us to a new view of how cells can convert from one phenotype to another. From these findings, we may begin to infer the adaptive capacity of a group of genes or cellular population given environmental challenges.

Mass arrest of major features of a cellular phenotype, such as cell cycle, transcription, and protein synthesis, can be accomplished using either drug treatment or transgenic knockdowns. Drug treatments have been done on selected cell lines to assess the time course of mRNA decay after exposure to a drug stimulus. It should be mentioned that any given drug treatment does not result in a uniform biochemical response neither across genes within a single cell population or between cell populations. However, a generalized effect of drug treatments on gene expression can still be observed. In previous studies this effect has proven to be a mosaic, as genes with different functions respond differentially to the drug stimulus [1, 2]. Knockdowns of specific genes have also resulted in shutting down expression using a transgenic construct and measuring the remaining mRNA [3]. While knockdown studies are focused on specific genes, techniques associated with mRNA decay allow for examining the expression of many genes in the context of a common stimulus. The benefit of using an mRNA decay approach involves being able to capture the behavior of a single gene’s expression against a background of widespread functional decay.

1. Dressaire, C. et.al (2009). Translatome and proteome exploration to model translation efficiency and protein stability in *Lactococcus lactis*. PLoS Comput Biol, 5(12), e1000606.
2. Perez-Ortin, J.E. (2008). Genomics of mRNA turnover. Brief Funct Genomics, 6(4), 282-291.
3. Archer, S. et.al (2008). Trypanosomes as a model to investigate mRNA decay pathways. Methods Enzymol, Volume 448, 359-377.

## C_t_ measurement

In order to measure the effect each drug stimulus has on mRNA molecule abundance, the C_t_ measurement was used as a heuristic for fluctuations that occur due to decisions made by cell populations. The raw C_t_ (or cycle threshold) measurement is defined as the number of cycles in a quantitative PCR reaction required to achieve a signal that is distinguishable from background noise [1]. The signal itself is a florescent detector that is inversely related to the presence of the target nucleic acid [2]. Therefore, a larger number of target molecules results in quicker detection by the florescence probe and a relatively low C_t_ value.

The Ct measurement can be converted to a relative measurement through the use of comparisons to a control. Depending upon analytical need, our Ct measurements were normalized in two ways: using a housekeeping gene (GAPDH) and using an untreated control. In this way, we were able to provide relative measurements that were independent of potential biological function and/or experimental effects.

1. Life Technologies (2011). Real-time PCR: understanding Ct. Application Note, Applied Biosystems.
2. Sigma-Aldrich (2008). qPCR Technical Guide. Sigma Life Science.

## Selection of Candidate Genes

In order to test our combination of a relatively obscure technique and systems model, we used a candidate gene approach to investigate expression across a pilot study and two experiments. An initial (non-quantitative) pilot experiment was used to validate the differential effects of our chosen genes (using standard PCR), and involved comparing HeK-293, early passage YFP-L10A fibroblasts, and late passage/senescent YFP-L10A fibroblasts. The choice of our candidate genes were based on genome annotation data and prior literature [1, 2]. While a high-throughput dataset would have potentially given us a more conclusive result, using genes representative of specific cellular identities and processes allowed us to take a first step in establishing our approach. Our candidates included genes that are thought to be highly regulated in many fibroblast lines (e.g. fibroblast-specific genes). As with the fibroblast-specific genes, our criterion included genome annotation and prior literature [1, 2]. Once the basic techniques were validated with our first experiment, we conducted a second experiment to extend these results by investigating the potential for detecting unexpected regulatory events using genes with no known function in fibroblasts. Unlike in the first experiment, only the SAP form of mechanism alteration was investigated. This was done because such unexpected regulatory events should explicitly involve upregulation of TLT. This can also serve as a test of false-positives for our polysome recovery techniques.

1. Satish, L. et.al (2008). Identification of differentially expressed genes in fibroblasts derived from patients with Dupuytren’s Contracture. BMC Med Genomics, 1, 10.
2. Abiko, Y., Hiratsuka, K., Kiyama-Kishikawa, M., Ohta, M., and Sasahara, H. (2004). Profiling of differentially expressed genes in human gingival epithelial cells and fibroblasts by DNA microarray. J Oral Sci, 46(1), 19-24

## Outline for Analysis

In order to validate TLT technique and then generalize it to regulatory dynamics, we will proceed through seven steps that will allow us to validate and characterize our technique for isolating TLT and modeling a direct comparison with TST. The analysis consists of two parts: validating our approach (e.g. measuring TLT and using the mechanism alteration approach) and the implementation of our systems model. To validate our approach, we will use fibroblast-specific and non-specific genes normalized by a single gene known to be involved with housekeeping functions.

The first step will be to conduct an initial evaluation using an exploratory data analysis technique called a quartile-quartile (or Q-Q) plot of the outcome for comparing TLT and TST C_t_ values. The next step is to establish decay profiles to determine what should be expected at each time point for the TLT and TST, respectively. To explore these differences (e.g. gene-specific) further, we will conduct a regression analysis for each gene’s transcription-related and translation-related RNA. As an alternate means of investigating the correspondence between TLT and TST in a gene-independent manner, the correlation coefficients are calculated for selected gene and treatment day combinations. This will allow us to determine for which genes there was a high correspondence between TLT and TST, and how this relative correspondence compares across genes and phases of biochemical change. To make more meaningful comparisons between experimental treatments, TST/TLT, and genes, linear decay rates were subject to a false positive analysis and mapped to a bivariate space.

## RESULTS

### Cell culture experiments

The first step in our analysis is to establish that TLT and TST can indeed be directly compared. Figure 1 is an initial statistical comparison of TLT and TST for each drug treatment using a Q-Q plot. The C_t_ values were transformed to z-scores, rank ordered, and then plotted on a bivariate graph (see Methods). By doing this for all three experimental conditions, we can both make a direct comparison between TLT and TST in addition to making relative comparisons between experimental conditions. In the case of AD, there appears to be more variability along the the TST axis. By contrast, MMC treatments shift most of this variability to the TLT axis. SAP treatments reveal variability that is more evenly distributed between the TLT and TST. In addition, SAP treatments reveal a greater number of outliers (z-score above 2.0 or below −2.0) along both axes. For details on our outlier strategy, please see Methods.

**Figure 1.**
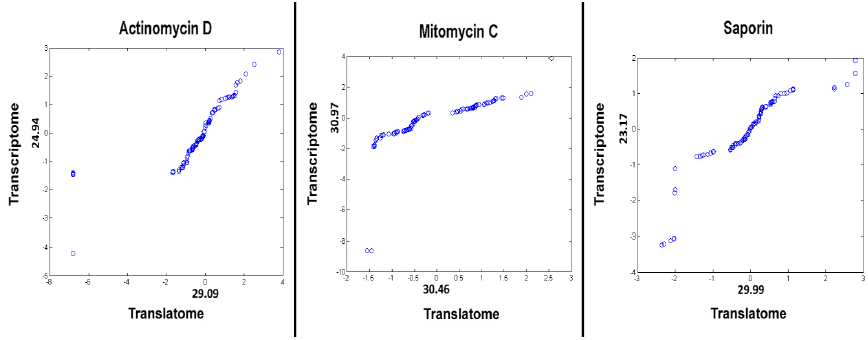
Q-Q plots for TLT (x-axis) vs. TST (y-axis) for all three drug treatments (units = z-score, Mean Ct value labeled along each axis at z-score of zero). Left: AD, Center: MMC, Right: SAP. NOTE: data not corrected for outliers.

Now that we have shown a basic difference between the two types of mRNA, we will now demonstrate that the C_t_ values sampled over time have complex dynamics that differ between TLT and TST in a gene-specific fashion. Aside from noisy gene expression, complex dynamics also result from information processing in cell populations given a particular stimulus at different points in time. These dynamics can be approximated using the mean values for each mRNA fraction /time point/gene combination. We have model this using a nonlinear curve-fitting model that accounts for mean fluctuations for each gene and mRNA fraction over time. In Figures, 2, 3, and 4, the profiles for MMC, AD, and SAP treatments respectively, are shown. The quantification of each treatment and normalized fraction of mRNA for 1d, 2d, and 3d were fit to a 2^nd^-order polynomial regression function. The most striking outcome is that there are increases in both TLT and TST at 2d and 3d. The increase at 2d in the TLT of AD-treated cells may be due to the aggregation of mRNA in the polysome. This may be a genome-wide survival strategy, as it is know that AD-treated cell cultures collapse by 8d post-treatment. Likewise, the aggregation of mRNA in MMC-treated cells may be related to transcription and translation of selected in response to the cell cycle blockade.

**Figure 2.**
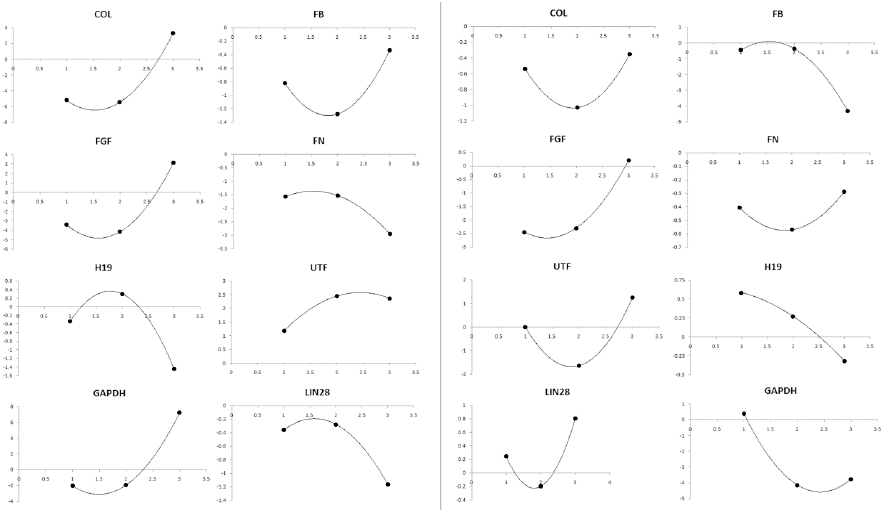
Decay curves (idealized using a 2nd-order polynomial) for MMC treatment. Time (days, x-axis) vs. mRNA quantification (y-axis). Left: TST. Right: TLT.

**Figure 3.**
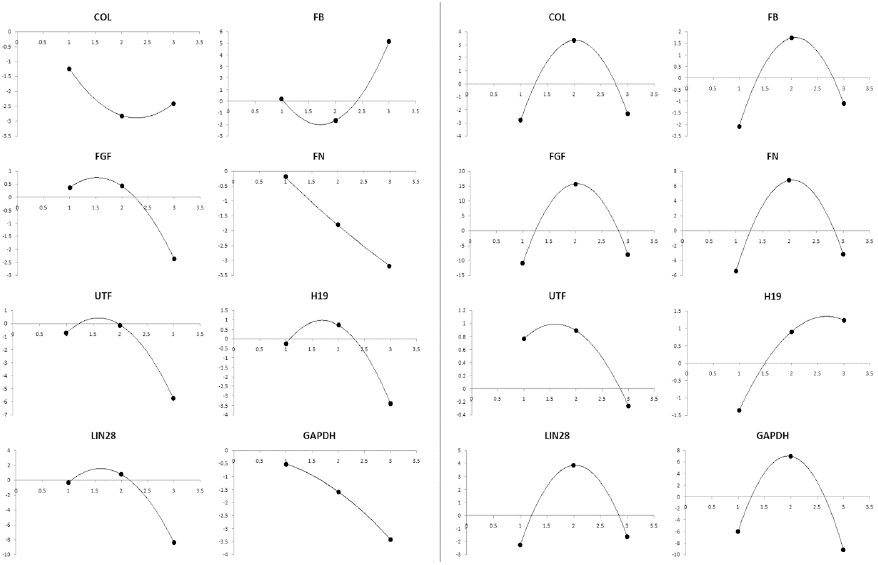
Decay curves (using a 2nd-order polynomial) for AD treatment. Time (days, x-axis) vs. mRNA quantification (y-axis). Left: TST. Right: TLT.

**Figure 4.**
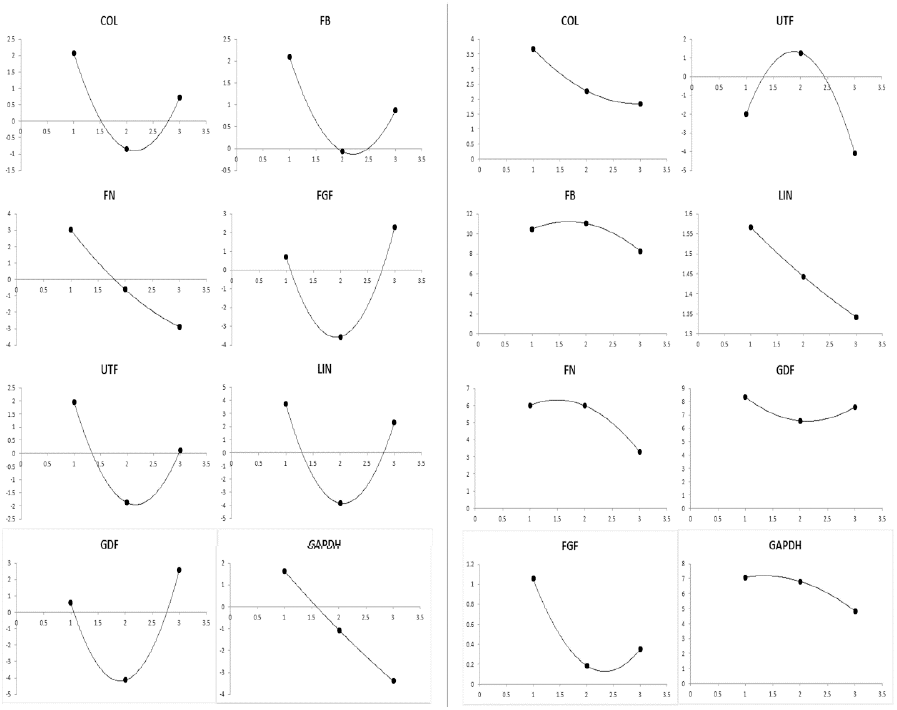
Decay curves (using a 2nd-order polynomial) for SAP treatment. Time (days, x-axis) vs. mRNA quantification (y-axis). Left: TST. Right: TLT.

Across several days, the mRNA profiles also appear to be gene specific (see results for MMC in Figure 2 and AD in Figure 3). A linear regression (Table 1) was used to calculate a decay rate in terms of C_t_ cycles per day. This represents the decay profile for each gene and treatment combination. If the response of a given gene is dominated by decay processes rather than dynamics, the linear regression should be highly significant. In Table 1 we find that roughly half (44%) of the gene and treatment combinations have significant decay components.

**Table 1.** Decay rate predicted using linear regression (units = cycles per day). Positive values (bold) represent decay, negative values (bold) represent aggregation.

|  | COL | FB | FN | FGF | UTF | LIN | GDF/H19 | GAPDH |
| --- | --- | --- | --- | --- | --- | --- | --- | --- |
| **SAP, TST** | **0.69** | **0.61** | **2.96** | **-0.81** | **0.92** | **0.70** | **-1.03** | **2.51** |
| p-value | 0.06 | 0.46 | 0.58 | 0.007 | 0.18 | 0.38 | 0.17 | 0.16 |
| **SAP, TLT** | **0.92** | **1.08** | **1.35** | **0.35** | **1.06** | **0.11** | **0.37** | **1.13** |
| p-value | 0.01 | 0.002 | 0.01 | 0.33 | 0.33 | 0.65 | 0.001 | 0.001 |
| **AD, TST** | **0.58** | **2.45** | **1.48** | **1.36** | **2.53** | **4.01** | **1.58** | **1.45** |
| p-value | 0.001 | 0.11 | 0.001 | 0.38 | 0.01 | 0.03 | 0.02 | 0.001 |
| **AD, TLT** | **-0.25** | **3.31** | **-1.15** | **-1.43** | **0.52** | **-0.33** | **-1.29** | **1.59** |
| p-value | 0.09 | 0.22 | 0.42 | 0.63 | 0.01 | 0.03 | 0.74 | 0.04 |
| **MMC, TST** | **-4.24** | **-0.24** | **0.69** | **-3.28** | **0.55** | **0.40** | **-0.59** | **-4.65** |
| p-value | 0.57 | 0.02 | 0.31 | 0.51 | 0.001 | 0.001 | 0.001 | 0.11 |
| **MMC, TLT** | **-0.09** | **1.93** | **-0.06** | **-1.32** | **-0.62** | **-0.28** | **0.45** | **2.08** |
| p-value | 0.60 | 0.06 | 0.24 | 0.17 | 0.94 | 0.003 | 0.91 | 0.001 |

Thus, not all gene and treatment combinations are dominated by decay. For the MMC treatment, we observe selective aggregation in both TLT and TST. The AD treatments exhibit more aggregation in TLT when compared to the corresponding TST. Meanwhile, the SAP treatments exhibit almost universal decay in both TLT and TST. This pattern of aggregation at 2d was confirmed in a second experiment (Figure 5). In this experiment, aggregation was also observed at 2d for selected specific and non-specific genes. This occurred in spite of massive cell death at 3d, which was significant enough to prevent cells from being assayed for analysis. This massive cell death leads to the assumption that any mRNA recovered at 3d would be greatly downregulated, thus replicating the curves shown in Figure 4.

**Figure 5.**
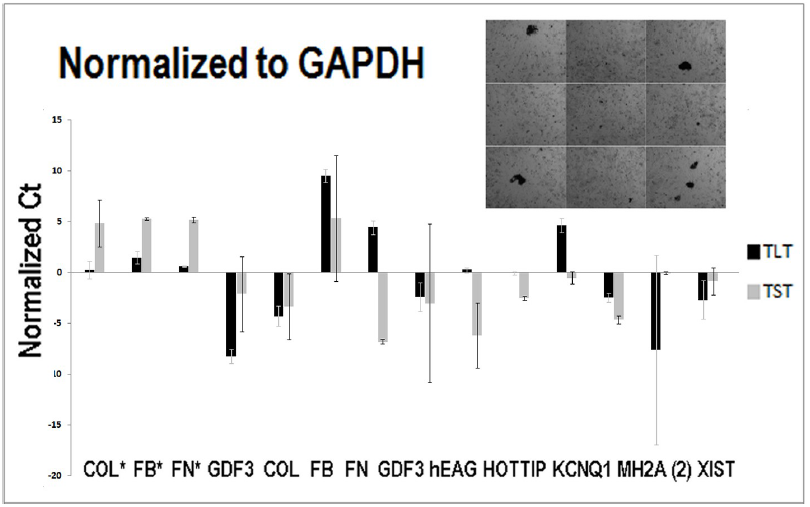
Confirmatory comparison of TLT (black) and TST (gray) for SAP treatment. Starred conditions (COL, FB, FN) assayed at 0d (control); all other conditions assayed at 2d post-treatment. All genes normalized to GAPDH. Upper right: microscopy demonstrating the degraded condition of fibroblasts at 2d post-treatment.

To further underscore the gene-specific relationship between TLT and TST, the correlation coefficients between these two fractions of mRNA are shown for selected genes across all treatments. The data were stratified by time point (e.g. 1d, 2d, and 3d). These results are shown in Table 2. The results demonstrate three distinct results. A moderate to high negative correlation for COL (-0.68), FN (-0.79), and GADPH (-0.76) occurs at 1d. A moderate to high positive correlation among fibroblast specific genes (COL, 0.67; FB, 0.76; FN, 0.59) as well as FGF (0.64) and GAPDH (0.62) occurs at 2d. Finally, a low to moderate correlation coefficient occurs for all selected genes at 3d. Breaking the analysis down in this way demonstrates the semi-independence of each mRNA fraction. In the case of housekeeping (e.g. GAPDH) and fibroblast-specific genes (e.g. COL, FB, FN), a correlation between TLT and TST indicates a stable maintenance of expression in the face of functional (or mechanism) disruption. The moderately positive correlation between TLT and TST of GAPDH at 3d (as opposed to the weak positive to negative correlation for all other genes tested) demonstrates this stability mechanism. In this case of GAPDH, TLT and TST seem to be strongly decoupled processes during 1d but loosely coupled during 2d and 3d. Overall, the results of our correlation analysis along with our nonlinear curve-fitting exercise suggest a generalized response at 2d that holds across replicates and experimental treatments.

**Table 2.** Statistical correlation (using correlation coefficient) between TLT and TST for each gene across all treatments and time points. NOTE: H19 and GDF were excluded from analysis.

|  | 1d | 2d | 3d |
| --- | --- | --- | --- |
| **COL** | -0.68 | 0.67 | 0.20 |
| **FB** | 0.33 | 0.76 | -0.45 |
| **FN** | -0.79 | 0.59 | 0.11 |
| **FGF** | -0.10 | 0.64 | 0.23 |
| **LIN** | -0.06 | -0.30 | -0.23 |
| **UTF** | 0.58 | -0.11 | -0.15 |
| **GAPDH** | -0.76 | 0.62 | 0.41 |

For purposes of understanding to what extent mRNA dynamics unfold in TLT and TST as parallel processes, linear decay rates (serving as the expected rate of degradation and calculated independently of the linear regressions) are shown in Table 1 for TLT and TST were mapped to a bivariate space for further inspection (see Figure 6). This space was divided into four quadrants: 1) aggregating TST, decaying TLT, 2) decaying TST, decaying TLT, 3) aggregating TST, aggregating TLT, and 4) decaying TST, aggregating TLT. Clusters were detected using a Euclidean distance metric. This transformation confirmed that all drug treatments yielded a generalized downregulating effect, as only 2 of the 24 datapoints reside in our aggregating TST, aggregating TLT quadrant. A nonparametric statistical test also reveals the number of points in our decaying TST, decaying TLT quadrant is 17% above random expectation (Figure 7).

**Figure 6.**
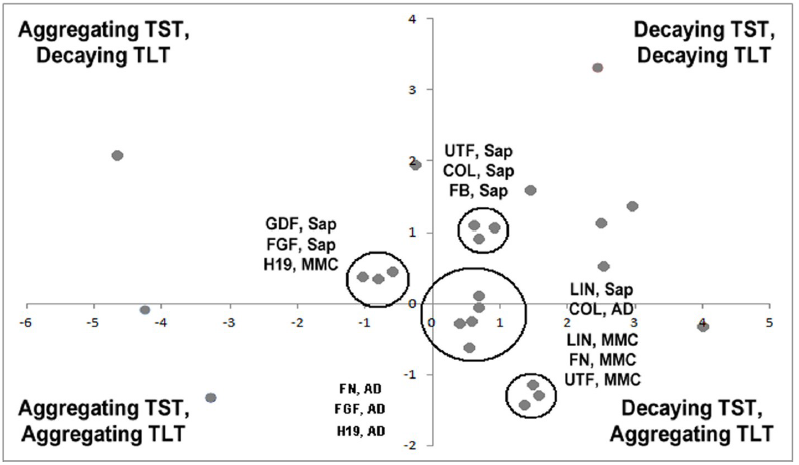
Linear approximation of decay (cycles per day) comparing TST vs. TLT for all drug treatments combined. Clusters (no statistical support, determined by Euclidean distance) shown within black circles. Abbreviations: SAP = Saporin, AD = Actinomycin D, MMC = Mitomycin C.

**Figure 7.**
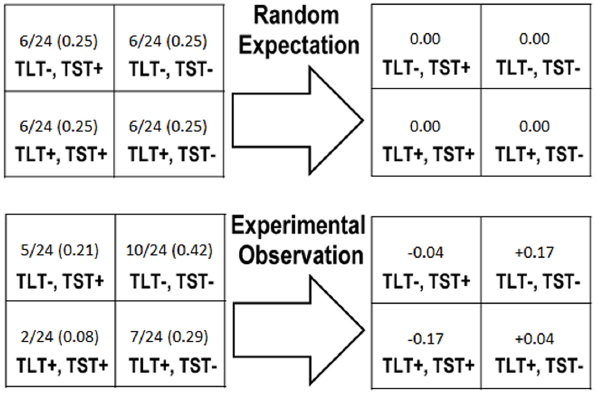
Nonparametric test demonstrating how much mRNA decay signatures deviate from random expectation. Lower left: fraction of samples observed in each quadrant. Lower right: deviation from expected value (example shown in table at upper left).Abbreviations: TST+,TLT-= aggregating TST, decaying TLT; TST-,TLT-= decaying TST, decaying TLT; TST+,TLT-= aggregating TST, aggregating TLT; TST-,TLT+ = decaying TST, aggregating TLT.

**Figure 8.**
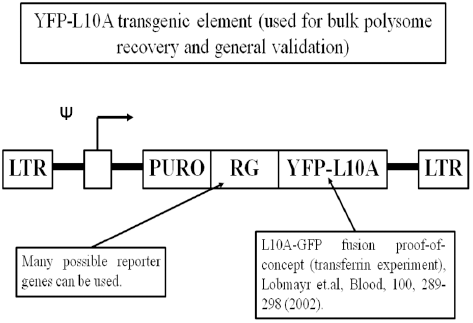
Example of the YFP-L10A construct used to validate polysome recovery. Reporter gene can be used in various contexts. A florescent YFP signal (no reporter gene) was used to identify cells carrying construct and to select mRNA.

This bivariate space also yielded four clusters. The first cluster contains three genes (UTF, COL, FB) under SAP treatment. All points for this cluster are in the decaying TST, decaying TLT quadrant. A second quadrant contains three genes. Two of these (GDF, FGF) are under SAP treatment, while the other (H19) is under MMC treatment. All points in this cluster were in the aggregating TST, decaying TLT quadrant. A third cluster contained three genes (FN, FGF, H19), all under AD treatment and residing in the decaying TST, aggregating TLT quadrant. The fourth cluster contains five instances of four genes (LIN, COL, FN, UTF), from all drug treatments. Three of these instances (LIN, FN, UTF) were a product of MMC treatment. The cluster straddles the boundary of the decaying TST, decaying TLT and the decaying TST, aggregating TLT quadrants, with four of the points contained in the latter.

## DISCUSSION

### Broader Technical Implications

There are several contemporary examples of how mRNA dynamics are studied that place our results in context. Barenco et.al [1] has examined mRNA dynamics by examining the DNA-damage response in the MOLT4 pathway, which acts in a cell-line and context specific manner. Using this model, they are able to use a hidden variable dynamical model to partition out variance associated with decay and other signatures over time from time-series microarray data. By contrast, our model does not explicitly partition variance related to different cellular processes. However, in holding certain mechanisms constant, we can gain insight into adaptive responses on a longer time-scale than do Barenco et.al [1] or Sharova et.al [2]. In this way, we can observe several potential nonlinear control mechanisms. What we do not know is whether these signatures are caused by previous gene expression patterns, a generalized adaptive response, or if they are an indicator of pure decay.

Rabani et.al [3] test the varying degradation hypothesis, which posits that changes in mRNA level over time are strongly affected by changes in degradation rate. This can be characterized by either a single or continuous shift in mRNA profiles, which tend to be gene specific. Likewise, we observe this gene-independent activity, which can be characterized either by a single shift (understood through nonlinear curve fitting) or a continuous shift (consistent with linear aggregation or decay response). These authors also use a technique called *4sU labeling* to separate newly transcribed mRNA from the total pool [1]. In doing so, it appears that the processing of mRNA at the site of transcription plays a role in shaping longer-term temporal functions. This is also consistent with the impulse model of Chechik and Koller [4], who suggest that mRNA dynamics are characterized by an abrupt early response coupled to a later transition towards a steady state. This regulatory output may be due to the pre-mRNA processing observed by Rabani et.al [3] or ribosomal functions [4]. This is also consistent with the complex responses observed in our data, particularly across different functional classes.

1. Barenco, M. et.al (2009). Dissection of a complex transcriptional response using genome-wide transcriptional modelling. Mol Sys Biol, 5, 327.
2. Sharova, L.V. et.al (2009). Database for mRNA half-life of 19,977 genes obtained by DNA microarray analysis of pluripotent and differentiating mouse embryonic stem cells. DNA Res, 16(1), 45-58.
3. Rabani, M. et.al (2011). Metabolic labeling of RNA uncovers principles of RNA production and degradation dynamics in mammalian cells. Nat Biotech, 29(5), 436-442.
4. Chechik, G. and Koller, D. (2009). Timing of gene expression responses to environmental changes. J Computational Biol, 16, 279–290.

## METHODS

### Drug treatment procedure and cell biology

Cells are plated in two (2) 6-well plates, and grown to 90% confluence. Cells are then treated using a pre-determined concentration of drug compound (MMC, 10mg/μL; AD, 50mg/μL; SAP, 20mg/μL). Three (3) wells per condition were treated. Treated cells are exposed to the drug compound for 24h. The drug compound is then removed and replaced with MEF culture medium. Samples are then harvested at 1d, 2d, and 3d post-treatment. Two control samples (3 wells each, untreated cells of the same type) per drug treatment were harvested at 100% confluence. Both control samples are quantified and averaged together to normalize (in a subtractive manner) 1d, 2d, and 3d samples.

### Cell culture

Cell culture work was done in the Cellular Reprogramming Laboratory. Transgenic human fibroblast lines (L10A-GFP/YFP) are used to conduct the experiments. The L10A-GFP/YFP elements are used as indicators of cellular state and gene expression. Please see Supplemental Information for further detail.

### Molecular biology protocols

Molecular biology work is done using PCR (primer validation) and qRT-PCR (quantification) techniques, while precipitation of RNA, TLT RNA, and polyribosomal fractions of cell lysate are done according to an established protocol. A set of primers that detect a representative assortment of genes are used to evaluate the various conditions and differences between TLT and TST fractions.

### TRAP protocol

Cultured cells are harvested by disruption with PBS at 4°C, trypsinization, and centrifuged to a pellet. To extract TLT, the pellet is suspended in polyribosome extraction buffer (PEB) and lysed via homogenization (see below). The lysate is then centrifuged at 2000g for 20 minutes at 4°C. The supernatant is then extracted and incubated with 10µL Triton X-100 and 7mg polar lipids (Avanti, Bessemer, AL) at 4°C for 30 minutes. Each tube is mixed via inversion before incubation. 30uL of anti-GFP magnetic beads (Miltenyi Biotech, Bergisch Gladbach, Germany) are then added to each tube, mixed via inversion, and then incubated end-over-end on a rotating tilt-table for 30 minutes at 4°C. Magnetic-assisted cell sorting (see below) are used to isolate the GFP^+^ mRNA molecules. mRNA in the resulting fraction are then isolated using the 3M Sodium Acetate protocol (see below). Once the aqueous phase is extracted, 1µL of Glycoblue per 100µL total volume is added to precipitate the maximum amount of RNA. The Glycoblue mixture is incubated for 10 minutes at −80°C, centrifuged at 14,000g for 25 minutes at 4°C, and then washed with EtOH. The resulting RNA pellet is resuspended in 20µL ddH_2_0 and cleaned using a Qiagen kit (see below).

### Polyribosome extraction buffer (PEB)

One volume of PEB is made by suspending 1.19g of HEPES, 5.60g of KCl, 0.51g of MgCl_2_, 0.4g of DTT, and 50µg of cycloheximide into 50mL of PBS. Four protease inhibitor tablets (Complete Mini, EDTA-free, Roche Diagnostics) are then dissolved into the suspension.

### Magnetic-assisted Cell Sorting (MACS)

MACS (Miltenyi Biotech, Bergisch Gladbach, Germany) is used to isolate the florescent reporter (GFP) from the lysate and recover all associated mRNA molecules. The lysate, which contains mRNA molecules tagged with Anti-GFP magnetic beads, are pipetted into columns exposed to a strong magnetic field. Once the flow-through is completed, the positive fraction is trapped in the column, and later washed out with buffer. All mRNA in the positive fraction of cells recovered from the column is associated with the L10A ribosomal moiety.

### Amazonia Database

The cell type specificity of primer amplification is determined by searching the Amazonia database of microarray results (http://amazonia.eu).

### Datasets and cell lines

For our pilot experiments, ADF human fibroblasts (obtained from gingival explants under Michigan State University Biomedical and Health IRB-approved protocols) are transfected with the YFP-L10A element (hereafter referred to as YFP-L10A fibroblasts) and HeK-293 (human cells commercially obtained from ATCC) cells were used. The ADF cell line has been obtained from a single individual, who has provided written informed consent. The ADF cell line has also been used in a previously published study [1]. The first experiment, which features all three mechanism alteration treatments, YFP-L10A fibroblasts are used. For the second experiment, featuring a replication of the SAP mechanism alteration treatment, YFP-L10A fibroblasts are also used. All cell lines are cultured in a standard fibroblast medium (DMEM+, FBS, L-glutamine, NEAA, antimycotic/antibiotic).

All data handling, statistical analysis, and computational modeling are done using R and Excel. The analyses involved four components: basic quantification of RNA, curve-fitting and regression analyses. Dataset is located on the Figshare repository at http://dx.doi.org/10.6084/m9.figshare.689894.

1. Suhr, S.T., Chang, E.A., Rodriguez, R.M., Wang, K., Ross, P.J., Beyhan, Z., Murthy, S., and Cibelli, J.B. (2009). Telomere Dynamics in Human Cells Reprogrammed to Pluripotency. PLoS One, 4(12), e8124.

### Primer Sets

Eight primer sets are used to assay cell-type specificity and relative expression in the proof-of-principle experiments for the TRAP method. Primers are designed using the NCBI primer design tool (http://www.ncbi.nlm.nih.gov/tools/primer-blast/). Collagen-1A2 (Forward, 5’-ATGGGCTTCGTGCCCAGTGC-3’; Reverse, 3’-CACAGCGGAACAGGCCAGGG=5’), Fibulin-5 (Forward, 5’-CAAGCCACGACCCGCTACCC-3’; Reverse, 3’-GGGCCCTTTGATGGGGCGTG-5’), Fibrillin-1 (Forward, 5’-CCTGGTGCTGCTGGACGGAC-3’; Reverse, 3’-CCACGAGGACCACGAAGCCC-5’), Growth Differentiation Factor 3 (Forward, 5’-TGGCTTTGGGCCAGGCAGTC-3’; Reverse, 3’-GGGAGACCCCAGTGGTCGCT-5’), Undifferentiated Transcription Factor (UTF) 1 (Forward, 5’-GGAACTCGGGTTGCCGGTGTC-3’; Reverse, 3’-GAGCTTCCGGATCTGCTCGTCGAAGG-5’), LIN-28 (Forward, 5’-TGCACCAGAGTAAGCTGCAC-3’; Reverse, 3’-ACGGATGGATTCCAGACCC-5’), Fibroblast Growth Factor (FGF) 4 (Forward: 5’-CCCACTGCACCCAACGGCACGC-3’; Reverse, 3’-TCATGGCCACGAAGAACCGGCTGGCCAC-5’), GAPDH (Forward, 5’-GCGGTCCCCCAGGTGAGAGT-3’; Reverse, 3’-GCAGTGCCCACAGCACCAAC-5’).

### Treatment Details

#### Mitomycin C treatment

Mitomycin C (MMC) is used to block cell cycle in treated cells, but not necessarily protein synthesis, the production of intracellular factors, or other functions of a cell. In at least one case, it has been found that cells can differentiate after MMC treatment [1]. Overall, the arrest of cell division can have a rate-limiting effect (e.g. decay) on mRNA and protein synthesis. However, it is not clear whether or not this signal will exhibit linearity.

#### Actinomycin D treatment

Actinomycin D (AD) has a more direct effect on blocking protein synthesis in treated cells, and explicitly targets the suspension of transcription and by extension RNA synthesis. The effect of treatment should be one of universal decay, with a cleaner signal when compared to MMC. However, there should be some residual effect from stabilizing factors such as microRNAs [2]. The kinetics of translation, however, is decoupled from translation in a way that allows us to observe a differential effect.

#### Saporin treatment

Cells are exposed to Saporin (SAP) for purposes of selectively targeting the destruction of polyribosomes, which may interfere with the docking and translation of polysome-associated mRNA [3]. Therefore, SAP is used to disrupt translation and retard aggregation of mRNA at the ribosome. This should result in greater decay of TLT when compared to TST. SAP treatment should have a similar effect on TLT that AD has on TST.

1. Filoni, S., et.al (1995). The inhibition of cell proliferation by mitomycin C does not prevent transdifferentiation of outer cornea into lens in larval *Xenopus laevis*. Differentiation, 58(3), 195-203.
2. Winter, J. and Diederichs, S. (2011). Argonaute proteins regulate microRNA stability: Increased micro RNA abundance by Argonaute proteins is due to microRNA stabilization. RNA Biol, 8(6), 1-9.
3. Bagga, S., Seth, D., and Batra, J.K. (2003). The Cytotoxic Activity of Ribosome-inactivating Protein Saporin-6 Is Attributed to Its rRNA N-Glycosidase and Internucleosomal DNA Fragmentation Activities. J Biol Chem, 278(7), 4813-4820.

### Other Supporting Information

The translatome (TLT) protocol used in this paper (bulk polysome) has been validated using both a transgenic construct (TRAP-YFP-L10A) in skin fibroblasts and a Northern blot/non-quantitative PCR approach. To ensure that our based protocol selectively yields only (or mostly) TLT-associated mRNA, we can create clonal lines selected for expression of the YFP-L10A element (see Figure 8).

In conjunction with the use of GFP-antibody magnetic beads and MACS (magnetic-assisted cell sorting) technology, the YFP-L10A element acts to "TRAP" RNA associated with the polysome. Subsequent detection with Southern blot techniques demonstrates that TLT can indeed be isolated, albeit in lesser quantities than transcriptome (TST). The number of cycles required to obtain a signal for each gene corresponds to the qPCR results for both TRAP isolated mRNA and non-TRAP isolated TLT. The non-TRAP isolated TLT mRNA (or PEB only protocol shown in Figure 9, frame 4) is a bit less pure (determined via spectrophotometry) than TLT mRNA isolated by TRAP. Nevertheless, the resulting mRNA is still analyzable and experimentally replicable.

**Figure 9.**
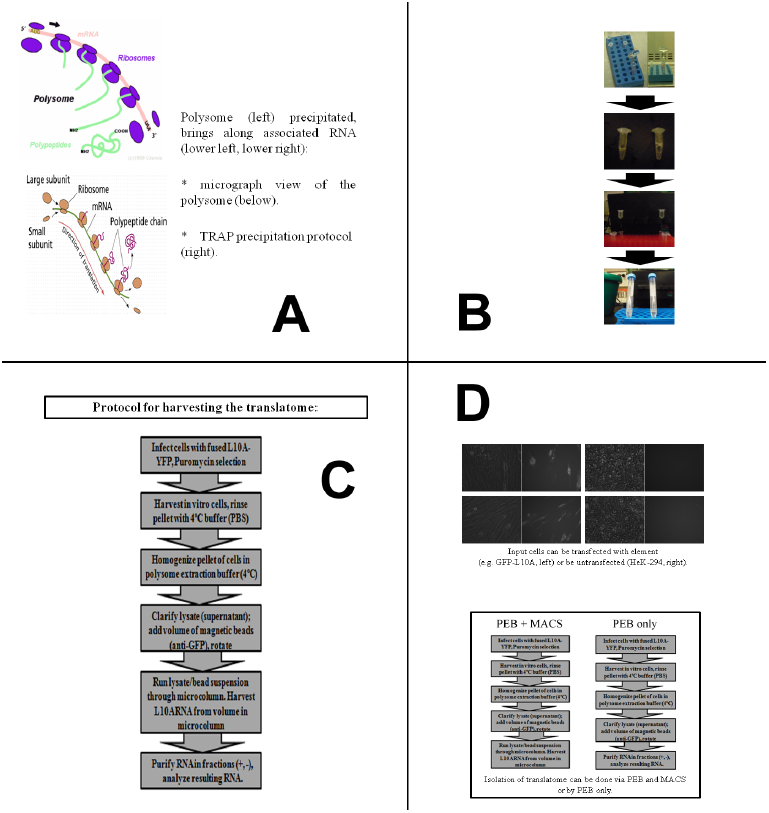
Image of direct polysome protocol process and L10A construct. A: basic biology of the polysome. B: process of polysome isolation and recovery using YFP-L10A element and magnetic beads. C: full protocol (step-by-step) for polysome isolation and recovery. D: steps involved in polysome recovery for both YFP-L10A cells (left) and regular cells (right). NOTE: the PEB only protocol does not require the addition of magnetic beads, but may be used as a control.

